# ErpY-like lipoprotein of *Leptospira* outsmart host complement regulation by acquiring complement regulators, activating alternate pathway, and intervening membrane attack complex

**DOI:** 10.1101/2021.05.27.446086

**Authors:** Saswat Hota, Md Saddam Hussain, Manish Kumar

**Affiliations:** Department of Biosciences and Bioengineering, Indian Institute of Technology Guwahati, Guwahati -781039, Assam, India

**Author notes:** corresponding author: Manish Kumar, Department of Biosciences and Bioengineering, Indian Institute of Technology Guwahati, Guwahati-781039, Assam, India.

**Keywords:** *Leptospira*, LIC11966, rErpY-like protein, the complement system

## Abstract

The survival of pathogenic *Leptospira* in the host pivots on its proficiency to circumvent the immune response. These pathogens evade the complement system in serum by enticing and amassing the serum complement regulators onto their surface. ErpY-like lipoprotein, a surface-exposed protein of *Leptospira* spp., is conserved and exclusively present in the pathogenic spirochete. The recombinant form of this protein is comprehended to interact with multiple extracellular matrix (ECM) components and serum proteins like soluble complement regulators factor H (FH) and factor I (FI). Here, we document that the supplementation of recombinant ErpY-like protein (40 µg/mL) in the host (humans) serum augments the viability of *E. coli* and saprophytic *L. biflexa* by more than 2-fold. Pure complement regulators FH and FI, when bound to rErpY-like protein, preserve their respective cofactor and protease activity mandated to cleave the complement component C3b. The supplementation of rErpY-like protein (40 µg/mL) in serum ensued in ∼90 % reduction of membrane attack complex (C5b-9/MAC) deposition through alternate complement pathway (AP) activation. However, rErpY-like protein could moderately reduce (∼16%) MAC deposition in serum through the classical pathway (CP). In addition, the rErpY-like protein solely activated the AP, suggesting its role in the rapid consumption and depletion of the complement components. Blocking the pathogenic *L. interrogans* surface with anti-rErpY resulted in an increase in MAC formation on the bacterial surface, indicating a specific role of the ErpY-like lipoprotein in complement-mediated immune evasion. This study underscores the role of the ErpY-like lipoprotein of *Leptospira* in complement evasion.

## Introduction

Pathogenic *Leptospira* spp. are the causative agents of leptospirosis, an emerging zoonotic disease of global importance (1). These are spirochetes that colonize the kidneys of rodents and can infect humans and domestic animals through soil or water contaminated with rodent urine (2). The workforce engaged in agriculture and livestock are the primary victims of the disease (3). The spirochetes invade the host through intact or injured mucous membranes then spread from the site of exposure to the target tissue through the circulatory system. Successful colonization in the host tissue counts on the pathogen’s ability to evade the host immune system, proliferate in blood, and adhere to and infiltrate host tissue (4). The complement system is the primary workhorse of the innate immune system against invading pathogens, including *Leptospira*. While the saprophytic leptospires are cleared upon coming in contact with the human serum, the pathogenic species of *Leptospira* have formulated multiple strategies to circumvent the complement system for continued survival in the host (5-8).

The human complement system encompasses over 50 proteins in the serum. Each complement protein is triggered in cascade-like steps, forming an effector complex known as membrane attack complex (MAC) that can perforate the target membrane (9). The complement activation occurs through the three major pathways: classical, lectin, and alternate (9). All these pathways converge on the cleavage of the central complement component, C3 (10). The complement activation through the classical pathway (CP) and lectin pathway (LP) banks on the availability of antigen-antibody immune complex on the target surface and the presence of distinct molecular patterns (carbohydrate moieties) on the target surface, respectively (11, 12). The alternative pathway (AP) is active in a spontaneous and nonselective way on all cell surfaces unless safeguarded by the regulatory proteins of the complement system. AP is activated when a small proportion of C3 molecules undergo a spontaneous conformational change in the blood, resulting in hydrolysis of the thioester group, forming an activated C3 called C3(H_2_O). A regulatory complement protein factor B (FB) binds C3(H_2_O), which is then cleaved by the circulating factor D (FD) to Bb and Ba, resulting in the formation of C3(H_2_O)Bb, the fluid phase C3 convertase of AP. Complement intermediate Bb is a serine protease that, in alliance with C3(H_2_O), can cleave more C3 to form C3b and C3a. C3b and Bb components bind covalently to nearby surfaces indiscriminately to form C3bBb, a membrane-bound C3 convertase. The events of recognizing the C3b by factor B, cleavage by factor D, and generation of more C3 convertase create a substantial amplification loop, which in some cases contributes up to 80% of the total complement response (13, 14). Exaggerated and indiscriminate complement activation on self-tissue has hazardous effects and can lead to many diseases (15, 16). To override excessive complement activation, several regulatory mechanisms consisting of soluble and membrane-bound regulators are in place to restrict complement activation against self-tissue (Schmidt, Lambris et al. 2016). The list of soluble negative regulators of the complement system that circulate in the blood is factor H (FH), factor I (FI), C4b binding protein (C4BP), C1 inhibitor (C1-INH), clusterin, vitronectin, and factor-H like protein 1 (FHL-1) (Schmidt, Lambris et al. 2016).

FH is a 155-kDa serum glycoprotein and embodies 20 homologous complement control protein (CCP) modules (17, 18). It acts as the central negative regulator of the AP in the free soluble phase and on the cell surface. FH competes with the Bb component of complement to bind with C3b. The displacement of the Bb component causes decay of surface-bound and fluid-phase C3 convertase (19-22). FH primary function is to act as a cofactor of the FI (a serine protease) to enable the cleavage of C3b into iC3b, an inactive variant (22-24). FH discerns distinct polyanions on the host cell surface as self-markers for complement regulation on host cells (25-27). The acquisition of FH by the pathogen also plays a leading role in inhibiting phagocytosis and promoting bacterial dissemination due to degradation of C3 fragments and generation of C3a and C5a anaphylatoxins (Merle, Church et al. 2015; Merle, Noe et al. 2015).

During the leptospiremic phase, the leptospires encounter the complement-mediated immune response through AP, nevertheless, can readily thwart its destruction (6, 7, 28-30). Pathogenic leptospires’ ability to strive against the alternative pathway has been reported for decades (6, 30). Millions of years of evolution have endowed leptospires along with many other pathogens to bind host FH through surface-exposed proteins, thus making use of the host regulatory system to escape the complement (8, 31-33). To date, multiple outer membrane proteins (OMPs) of pathogenic *Leptospira* have exhibited interaction with FH, including, LenA (LfhA, Lsa24), Len B, LcpA, LigA, LigB, Lsa23, and ErpY-like lipoprotein (34-40). Lately, we have reported the moonlighting effect of ErpY-like lipoprotein (LIC11966), a highly conserved pathogen-specific virulence factor with multiple functions in pathogenic *Leptospira* spp. The recombinant ErpY-like protein can acquire FH directly from human plasma *in vitro*, and to our belief, it is the only known bacterial protein that exhibits FI binding (36). This encouraged us to study the role of ErpY-like lipoprotein in the context of complement evasion.

This study reports that the rErpY-like protein acquires the complement regulators (FH and FI) in a catalytically active state and imposes an inhibitory effect on MAC deposition. The rErpY-like protein activates the alternative pathway, depletes the complement components, and in turn, impedes the complement-mediated clearing of nonpathogenic bacteria (*E. coli* and *L. biflexa*).

## Materials and methods

### Bacterial strains, culture media and, proteins

The saprophytic *L. biflexa* serovar Patoc strain Patoc 1 and the pathogenic *L. interrogans* serovar Copenhageni strain Fiocruz L1-130 were grown as described before (41). The *Escherichia coli* strain DH5α or BL21 (DE3) was grown at 37°C in Luria-Bertani (LB) broth medium (M1245; HiMedia) or LB agar (M1151; HiMedia) with or without ampicillin (61314; SRL) or kanamycin (99311; SRL) at a concentration of 100 µg/mL for expression of recombinant proteins. Recombinant proteins (ErpY-like protein, Loa22, and ClpP1) of *L. interrogans* with 6×His-tag at its N-terminal were overexpressed and purified as described previously (36, 41, 42). The commercially available complement factor H (FH; C5813; Sigma), factor I (FI; C5938; Sigma), C3b (204860; Sigma), C4BP (C4BPB-184H; Creative Biomart) were outsourced through local vendors.

### Immunoelectron microscopy

Immunoelectron microscopy of *L. interrogans* was performed to localize the ErpY-like lipoprotein, as described elsewhere, with slight modifications (43, 44). Spirochetes were washed twice in phosphate buffer saline (PBS) and fixed with 1% glutaraldehyde. Spirochetes were then resuspended in PBS for 30 min at 25°C. After two more washes in PBS, spirochetes were applied to electron microscopy grids (# AGS147-4H; Agar Scientific). Spirochetes were incubated for 60 min with rErpY-like antibodies or preimmune serum diluted 1:100 in PBS-T buffer. PBS-T buffer comprises PBS, 1.5% bovine serum albumin (BSA), and 0.05% Tween 20. After four washes with PBS containing 1% BSA, spirochetes were incubated for 60 min with goat anti-mouse gold-labeled antibody at 1:30 dilution (G7652; Sigma). The grid was further washed, and the spirochetes were stained with 2% uranyl acetate and were examined by field-emission transmission electron microscope (FETEM; JEOL-JEM-2100F) at an accelerating voltage of 80 kV.

### Serum susceptibility assay

The *E. coli* DH5α cells cultured in Luria Bertani (LB) broth were harvested at the exponential growth phase. The cells were washed once in PBS++ (137 mM NaCl, 2.7 mM KCl, 8 mM Na_2_HPO_4_, 2 mM KH_2_PO_4_, 1 mM MgCl_2_, 0.15 mM CaCl_2_) and adjusted to 4×10^4^ CFU/mL. Diluted normal human serum (NHS) (Sigma; H4522) in PBS++ (0.5%) was pre-incubated with different concentrations (10, 20 and 40 µg/mL) of rErpY-like protein or rClpP1 for 15 min at 37 °C. After that, approximately 2×10^3^ bacterial cells were incubated with the pre-mixed diluted serum (0.5%) for an hr at 37°C in a total volume of 100 µL. Following the incubation, the complement activation reaction was stopped by placing the tube on ice (1 min) and then the reaction mixtures were spread onto the LB-agar plates. The viable bacterial counts were determined by counting the *E. coli* colonies (CFU) after 14 h incubation at 37 °C. The results of viable *E. coli* (CFU count) were expressed in percentage after serum treatment, where the CFU count for untreated bacteria was considered 100%. For *L. biflexa* viability assay, cells were grown until the 4th day (in EMJH medium at 29 °C), harvested, washed once with PBS, and resuspended in EMJH after adjusting to 1×10^9^ cells/mL. Diluted NHS (40%) in EMJH was pre-incubated with the rErpY-like protein or rClpP1 at various (10, 20 and 40 µg/mL) concentrations for 15 min at 37 °C. After that, 5×10^7^ spirochete was incubated with the diluted NHS (40%) for an hour at 37 °C in a total volume of 100 µL. Following the incubation, the reaction mixture (100 µL) was inoculated in 1 mL of EMJH medium, and the spirochete was allowed to grow for 24 h at 29 °C. The viable leptospires at 24 h were counted on a Petroff-Hausser counting chamber under phase-contrast microscopy. For control, the complement component of NHS was heat-inactivated (ΔNHS) at 56°C for 30 min. The bacterial cells *E. coli* and *L. biflexa* were also incubated with 0.5% and 40%ΔNHS, respectively. The results of viable *L. biflexa* (cell count) are expressed in percentage after serum treatment, where the *L. biflexa* viable cell count for untreated spirochete was 100%. The Student’s two-tailed t-test was performed for statistical analysis.

### MAC deposition assay

The inhibitory effect of the rErpY-like protein on MAC (C5b-9) deposition was assessed *in vitro* as described previously (38, 45), with modifications. Zymosan A (Z4250; Sigma) was solubilized in 0.15 M sodium chloride at a concentration of 10 mg/mL. Zymosan was activated by placing it in a boiling water bath for an hour and then centrifuged for 30 min at 4000 g. The supernatant obtained was discarded, and the zymosan A residue was suspended evenly in the PBS to the desired concentration of 10 mg/mL as described elsewhere (46). To aggregate human IgG (I4506; Sigma), its solution (2 mg/mL in PBS) was incubated at 56°C for 15 min (47, 48). The microtiter plate was coated with activated zymosan A (20 µg/mL, for alternative pathway) or aggregated human IgG (2.5 µg/mL, for classical pathway) by incubating at 4°C for 16 h. NHS was diluted to 20% and 0.5% using the diluent GVB MgEGTA (#IBB-320X, Boston Bioproducts) and GVB++ (#IBB-300X), respectively. The two diluent selectively allow either the alternative (GVB MgEGTA) or classical pathway (GVB++) activation in NHS. The diluted NHS (20% and 0.5%) were pre-incubated with different proteins (rErpY like protein, rClpP1 or BSA) in increasing concentrations (10, 20 and 40 µg/mL) at 37 °C for 15 min. The individual reaction mixtures and the diluted NHS with no proteins were then added to a plate coated with activated zymosan A and aggregated human IgG. The plate was allowed to incubate at 37 °C for an hour. The deposited C5b9 complex in the reaction mixture was detected using rabbit anti-C5b-9 pAb at 1:1000 dilution (204903; Sigma) and HRP conjugated anti-rabbit pAb (1:5000) as primary and secondary antibodies, respectively. The secondary antibody binding was detected by adding 50 μL of tetramethylbenzidine (TMB) peroxidase substrate (cat# 34022, Thermo Fisher Scientific). The peroxidase reaction was terminated by adding 50 μL of H_2_SO_4_ (1 M) before the final optical density (OD) was taken using an ELISA plate reader (Multiskan Go, Thermo Scientific) at the wavelength of 450 nm. The OD value obtained for the reactions were converted to MAC deposition (%), considering the OD value obtained for unsupplemented NHS (control) as 100% and plotted. The Student’s two-tailed t-test was performed for statistical analysis.

### Complement activation assays

Activation of complement pathways in response to recombinant protein antigens was performed as described previously (45). Microtiter plates (cat# 941196, Tarsons) were coated either with 50 µL of rErpY-like protein (20 µg/mL), aggregated human IgG (2.5 µg/mL for the classical pathway), or zymosan (20 µg/mL for the alternative pathway) in PBS overnight at 4 °C. Wells coated with BSA (20 µg/mL) was used as a control. The plates were washed with PBS-T and were then blocked with BSA (3% in PBS) for 2 h at 37 °C. After blocking, the pre-diluted NHS in GVB++ (for classical) or GVB MgEGTA (for alternative pathway) was added to the plates and incubated for 20 minutes at 37 °C. Following the incubation, the deposited complement protein C3b was detected using the anti-C3b pAbs (1:1000 in PBST) for 2 h, and horseradish peroxidase-labeled secondary Abs against rabbit (1:5000 in PBST) for an hour at 37 °C. The absorbance values (450 nm) obtained for rErpY-like protein were compared with that of BSA (negative control) using the Student’s two-tailed t-test.

### Cofactor and protease activity assay

The microtiter plate was coated with 50 µL of rErpY-like protein (20 µg/mL) or rLoa22 (20 µg/mL) in PBS overnight at 4 °C. The plates were washed thrice with PBS-T and were blocked with BSA (3%) for 2 h at 37 °C. The plates were then overlaid with FH (2 µg) for 2 h at 37 °C. FI (0.25 µg) and C3b (0.5 µg) were added to the wells after rinsing off the unbound FH. The protease activity of FI bound to rErpY-like protein was assessed by overlaying FI (2 µg) on rErpY-like or rClpP1 coated wells for 2 h at 37°C followed by washing and addition of FH (0.5 µg) together with C3b (0.5 µg). To assess the ability of the antibody-blocked rErpY-like protein to capture FH or FI, mouse ErpY-like polyclonal antibodies (1:500) was added into microtiter plates coated with rErpY-like protein and then incubated for 2 h at 37°C. Unbound antibodies were washed away, and the wells were overlaid with either FH (2 µg) or FI (2 µg). Mixture of C3b (0.5 µg) + FI (0.25 µg) or C3b (0.5 µg) + FH (0.5 µg) were added and incubated with the FH or FI overlaid wells, respectively. All the reactions were performed for a duration of 4 h at 37°C. Reactions omitting FH were run as negative controls, and the reactions lacking the recombinant proteins were used as positive controls. The supernatant was resolved on 12% SDS polyacrylamide gel, and the cleavage fragments of C3b were detected by immunoblotting with a polyclonal anti-human C3b antibody (HPA020432; Sigma) (1:1000), and HRP conjugated anti-rabbit pAb (1:5000).

### MAC deposition on pathogenic spirochetes pretreated with rErpY-like antibodies

Five-day-old *L. interrogans* culture (1.3×10^8^ cells/mL) was harvested by centrifugation at 3000g. Spirochetes (1.3×10^8^) were treated with either mouse anti-rErpY-like or preimmune mouse serum (1:50) as the control for 1 h at 37 °C. The complement of the serum used was inactivated by incubating at 56 °C for 30 min before treating it with the spirochetes. The spirochetes pretreated with anti-sera and the untreated ones were incubated for 90 min at 37 °C, with 20 % NHS (diluted with GVB-MgEGTA) as a source of complement factors. Samples were then incubated on ice for 1 min to stop the complement activation. The cells were recovered by centrifugation, resuspended in PBS, and coated overnight onto the wells of a microplate at 4 °C. The deposition of the MAC on the spirochete surface was evaluated using anti-human C5b-9 antibody (1:1000) and HRP conjugated anti-rabbit antibodies (1:5000) as described above. The absorbance values obtained for MAC deposition on spirochetes blocked with anti-rErpY-like were compared statistically with untreated cells and those blocked with preimmune mouse serum (Student’s two-tailed t-test).

## Results

### ErpY like protein is an outer membrane surface-exposed protein on whole-cell immunogold electron microscopy

Using phase separation and proteinase K accessibility assay, our laboratory group had previously demonstrated the subcellular location ErpY-like lipoprotein to be an outer-membrane surface-exposed protein. The ErpY-like lipoprotein in *Leptospira* was annotated after the ErpY protein of *Borrelia*, showing 26% sequence identity (49). Borrelial ErpY comes under the OspF protein family (50). The proteins (ErpA, ErpC and ErpP) encoded by the borrelial *erp* (Osp**E**/F-**r**elated **p**rotein) (51) genes have been reported to act as complement regulator-acquiring surface proteins (CRASPs), implying their role in complement evasion (33, 52-57). In this study, immunoelectron microscopy further validated ErpY-like lipoprotein is a surface-exposed protein in pathogenic spirochete using gold-labeled secondary antibodies. The electron micrograph of pathogenic *L. interrogans* exhibited gold particles on their intact surface when incubated with specific serum against ErpY-like lipoprotein, unlike the ones incubated with the preimmune serum (Fig. 1). Whole-cell immunogold electron microscopy methods is advantageous, since the integrity of the outer membrane of the organism can be addressed from the images. Here the results of control are unambiguous and therefore the conclusions regarding surface-exposure are certainly valid. However, the disadvantage of this assay is that it may not reflect the total distribution of the ErpY-like lipoprotein in the organism. The expression of ErpY-like lipoprotein by pathogenic *Leptospira* spp. persuaded us to use these organisms to understand the role of ErpY-like lipoprotein in complement evasion.

**Figure 1.**
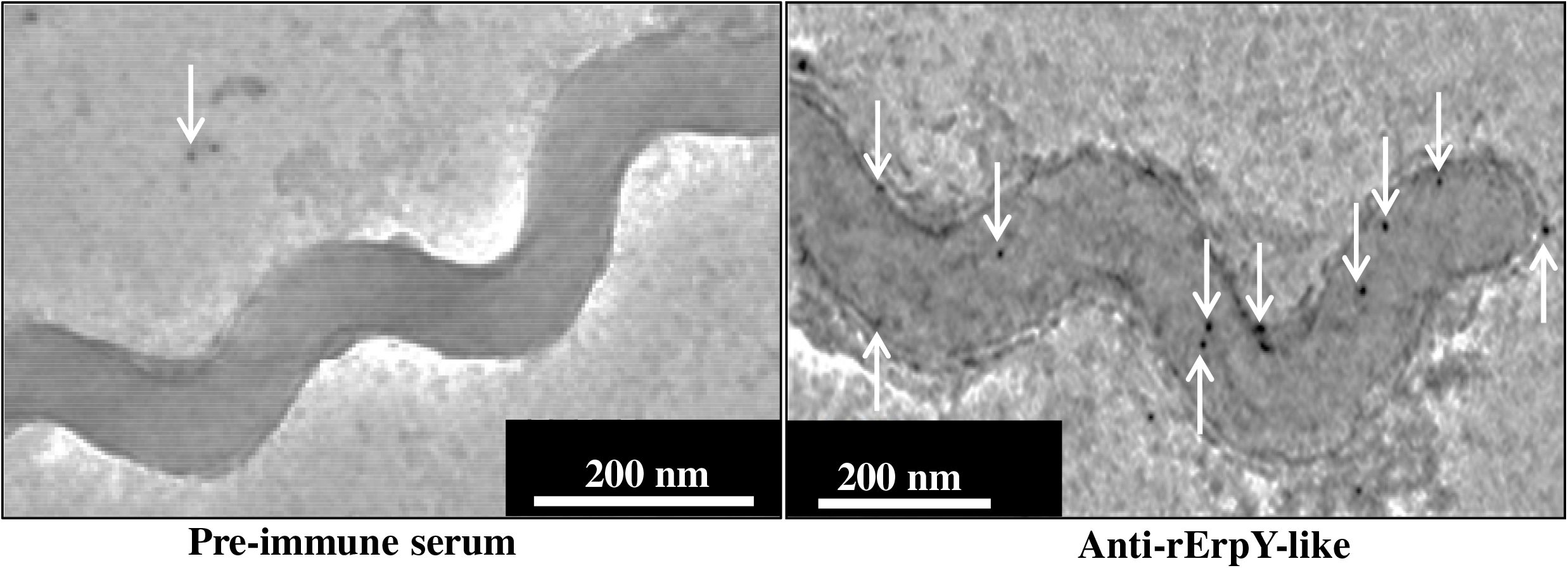
Whole cell electron micrograph of pathogenic *Leptospira* shows ErpY-like lipoprotein is an outer membrane surface-exposed protein. Pathogenic spirochetes were treated with rErpY-like protein antibodies (1:1000) or pre-immune serum (1:1000). Specific binding of antibodies was probed with the gold-conjugated anti-mouse secondary polyclonal IgG (1:20). The white arrows indicate the specific binding of anti-mouse IgG–colloidal-gold conjugates to the surface-exposed ErpY-like protein of spirochete.

### The presence of rErpY-like protein inhibits complement-mediated cytotoxicity of human serum on *E. coli* and *L. biflexa*

Pathogenic *Leptospira* mimics the host strategy to evade the toxicity caused by the complement activation in the blood by hijacking the host regulatory protein of complement activation like FH (5-8). Our laboratory had previously reported the ability of the ErpY-like lipoprotein of *L. interrogans* to acquire the complement regulator FH directly from human plasma (36). However, the role of acquiring the regulatory FH by rErpY-like protein from human plasma is not evident. Thus, it was intriguing to investigate if supplementation of rErpY-like protein in normal human serum (NHS) can enhance the viability of the *E. coli* and the model spirochete *L. biflexa*. The *L. biflexa* is a saprophytic spirochete that does not encode for ErpY-like lipoprotein in its genome (36). The susceptive *E. coli* cells, when was treated with NHS (0.5%) supplemented with rErpY-like protein (10 µg/mL) for an hour, a nearly two-fold rise in the viability of the cells was recorded versus the cells treated with only NHS (Fig. 2A). The measured percentage of viability (colony-forming unit, CFU count) of *E. coli* treated with only NHS was 37.7%. Interestingly, the viability increased to 71.5% when *E. coli* cells were treated in NHS supplemented with rErpY-like protein (10 µg/mL). Further increase of rErpY-like protein concentration (40 µg/mL) in NHS leads to augmentation in the viability of *E. coli* up to 88% (Fig. 2A).

**Figure 2.**
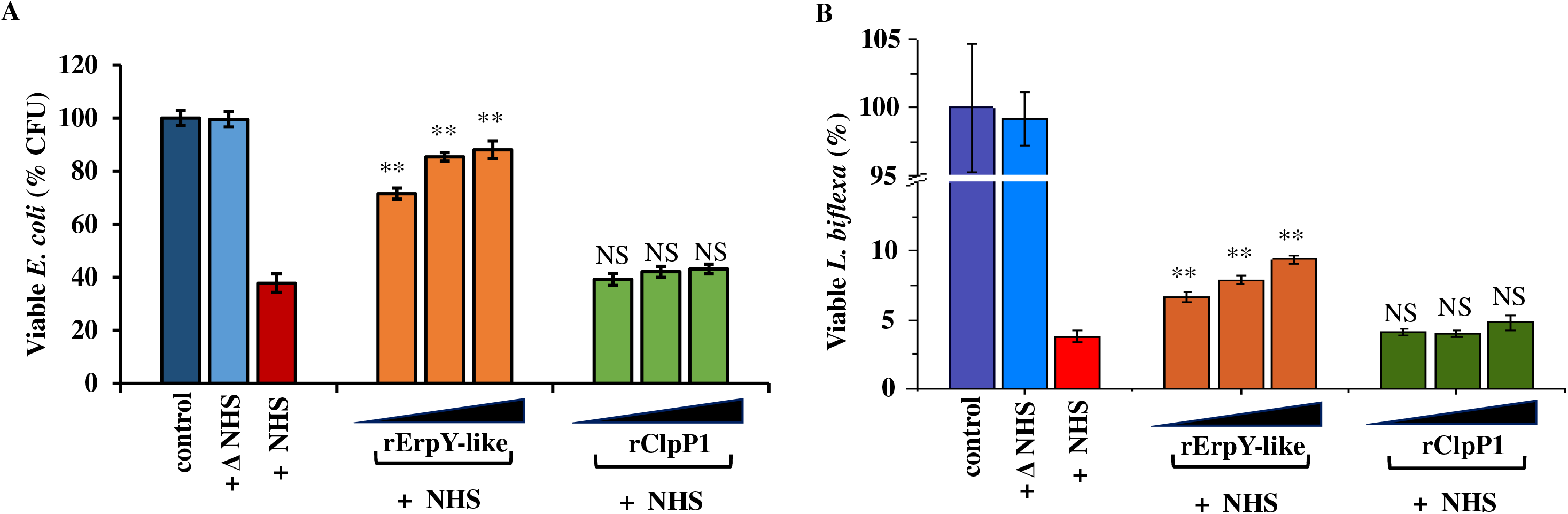
Recombinant ErpY-like protein increases the viability of *E. coli* and *L. biflexa* in normal human serum (NHS). **A**. Viability of *E. coli* in NHS supplemented with or without rErpY-like protein. Complement-mediated cytotoxicity on *E. coli* was reduced when treated in NHS (0.5%) supplemented with increasing concentration (10, 20 and 40 µg/mL) of rErpY-like protein. rClpP1 is a negative control used at the equivalent concentration for 1 h. The results of viable *E. coli* colony forming unit (CFU) count were expressed in percentage after serum treatment, where the CFU count for untreated bacteria (control) was considered 100%. Heat-inactivated NHS is denoted as ΔNHS. **B**. Viability of *L. biflexa* in NHS supplemented with or without rErpY-like protein. Recombinant ErpY-like protein reduces complement-mediated cytotoxicity on saprophytic *L. biflexa*. Saprophytic spirochetes were treated with NHS (40%) supplemented with or without rErpY-like protein in increasing concentrations (10, 20 and 40 µg/mL) for 1 h. The results of the viable leptospire count were expressed in percentage after serum treatment, where the cell count for untreated leptospires (control) was considered 100%. The assays were performed in three replicates, and the data shown are representative of three independent experiments. The error bars signify the standard error of the data sets. Statistical analyses were performed by the two-tailed Student’s t test where the viable CFU or the cell counts obtained for NHS treated bacteria (*E. coli* and *L. biflexa*) were compared with viability data of the control. (**, *p* < 0.001)

Similarly, *L. biflexa* treated with only NHS (40%) for an hour demonstrated 3.8% viability. On the other hand, the viability of the spirochetes improved to 6.6% when rErpY-like protein was supplemented in NHS. Thus, a nearly two-fold improvement in the viability was observed upon supplementation of rErpY-like protein (10 µg/mL) in NHS (Fig. 2B). The increase in spirochete viability (%) in the presence of ErpY-like protein was dose-dependent, similar to that of *E. coli*, where *L. biflexa* viability got improvised up to 9.4% (Fig. 2B). The rClpP1 and BSA was used as the control in the viability assay. The rClpP1 is a caseinolytic protease of pathogenic *Leptospira* that forms a supramolecule like rErpY-like protein (Dhara, Hussain et al. 2019). An equivalent amount of rClpP1 failed to impede the complement-mediated cytotoxicity in *E. coli* (Fig 2A) and saprophytic spirochete (Fig 2B), suggesting the role of rErpY-like protein to be specific in augmenting the viability.

### Regulatory complement factor (FH or FI) in a bound state with rErpY-like protein retain its C3b cleavage activity

FH is the primary negative regulator of the AP (Weiler, Daha et al. 1976; Whaley and Ruddy 1976; Whaley and Ruddy 1976). It forms an active complex with C3b, which is then acted upon by FI (a serine protease) to mediate C3b break down into an inactive form (iC3b) (58). It is understood that FI cleaves the C3b α’ chain (105 kDa) at Arg1303 and Arg1320 residues (59, 60). Proteolytic cleavage of C3b leads to the creation of iC3b molecule (173.6 kDa) comprising of the β chain (75 kDa), α’ (60.9 kDa), and α’ chain-2 (37.7 kDa) linked together by two disulfide linkages. In addition, another cleavage product, C3f (1.8 kDa), is released into the plasma. Active FI can further process the iC3b α’ (60.9 kDa) chain into C3c α’ chain-1 (22.5 kDa) and C3dg (38.39 kDa) by cleaving at the Arg954 residue (59). For clarity, the series of C3b cleavage episodes have been illustrated (Fig. 3).

**Figure 3.**
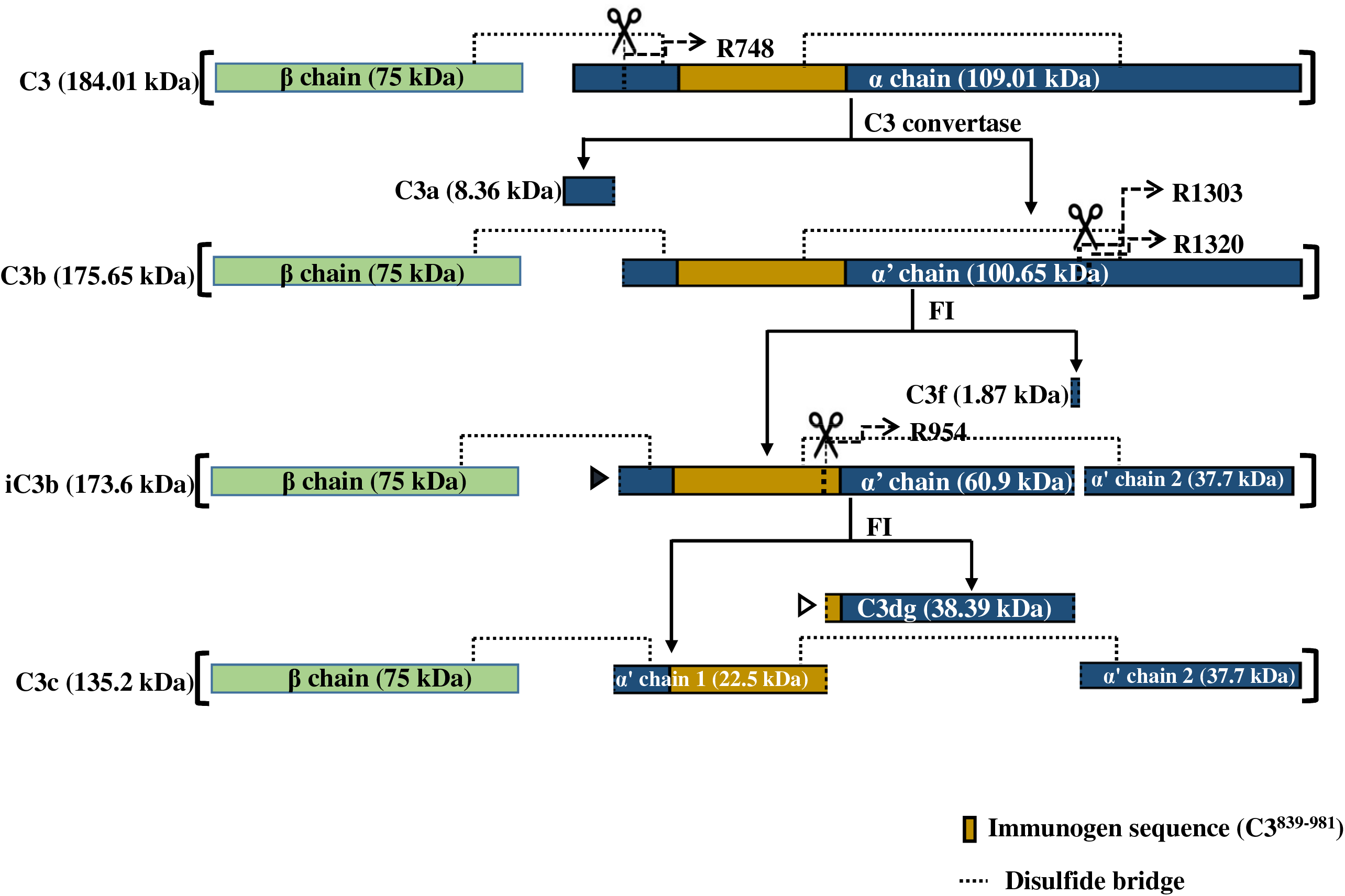
Illustration of complement factor C3 processing under physiological condition. C3 molecule (184 kDa) comprises a β chain (green) and an α chain (blue) linked through a disulfide bridge (dotted line). C3 convertase cleaves the α chain at Arg748 residue generating C3b (175.6 kDa) and C3a (8.3 kDa) fragments. FI cleaves at the Arg1303 and Arg1320 residues of the C3b α’ chain (100.6 kDa) in the first cleavage event producing the inactive C3b (iC3b; 173.6 kDa) and the C3f (1.8 kDa) fragments. FI consequently cleaves the iC3b α’ chain (60.9 kDa) at the Arg954 residue and generates the C3c (132.2 kDa) and the C3dg (38.3 kDa). The yellow segment of the α chain depicts the peptide sequence (C3^839-981^) that can be recognized by the antibody probe used in this study (rabbit anti-human C3b polyclonal antibodies). The α’ chain (60.9 kDa) and the C3dg (38.2 kDa) fragment detectable by the anti-C3b antibodies are indicated by the solid and the hollow triangles respectively.

FH acquisition, either in its active or inactive form by OMPs of pathogenic bacteria, is a known strategy to escape complement (61). In pathogenic *Leptospira*, multiple OMPs (LfhA, Len family protein, Lig family proteins, Lsa23) are reported to acquire the catalytically active host FH and mimic the host protective mechanism to evade the complement system (Verma, Hellwage et al. 2006; Stevenson, Choy et al. 2007; Castiblanco-Valencia, Fraga et al. 2012; Siqueira, Atzingen et al. 2013; da Silva, Miragaia Ldos et al. 2015; Siqueira, Atzingen et al. 2016). In contrast, PgtE of *Salmonella enterica* (62) and FhbB of *Treponema denticola* (63) deactivate FH upon binding, resulting in the depletion of complement constituents by unregulated activation of AP. The human serum susceptibility assays on *E. coli* and saprophytic spirochetes encouraged us to determine whether FH under bound state to the rErpY-like protein retains its cofactor activity or gets deactivated. Thus, a protease assay to cleave C3b was executed *in vitro* with FH on a microtiter plate, immobilized with or without rErpY-like protein. On immunoblot with C3b antibodies, two C3b fragments, α’ (60.9 kDa) and C3dg (38.39 kDa), were detected in the protease reaction containing FH and FI (Fig. 4A). It is to be noted that C3b antibodies (commercially outsourced) were generated against a specific epitope of C3 protein (C3^839-981^). The α’ fragment (60.9 kDa) was detected in the reaction where FH was preincubated with the immobilized rErpY-like protein. However, the C3dg (38.39 kDa) fragment was not detected in this reaction. As the cleavage product of C3b (α’ fragment, 60.9 kDa) is seen, it is evident that FH maintains its cofactor activity in the bound state to rErpY-like protein for the serine protease FI. In contrast, no such cleavage of the C3b α’ chain (105 kDa) was detected when FH was preincubated with the immobilized Loa22 (Fig. 4A). Similarly, the protease assay on C3b that lacked anyone regulatory complement protein (FH/FI) or both did not demonstrate any cleavage within C3b, implying the assay to be rErpY-like protein specific.

**Figure 4.**
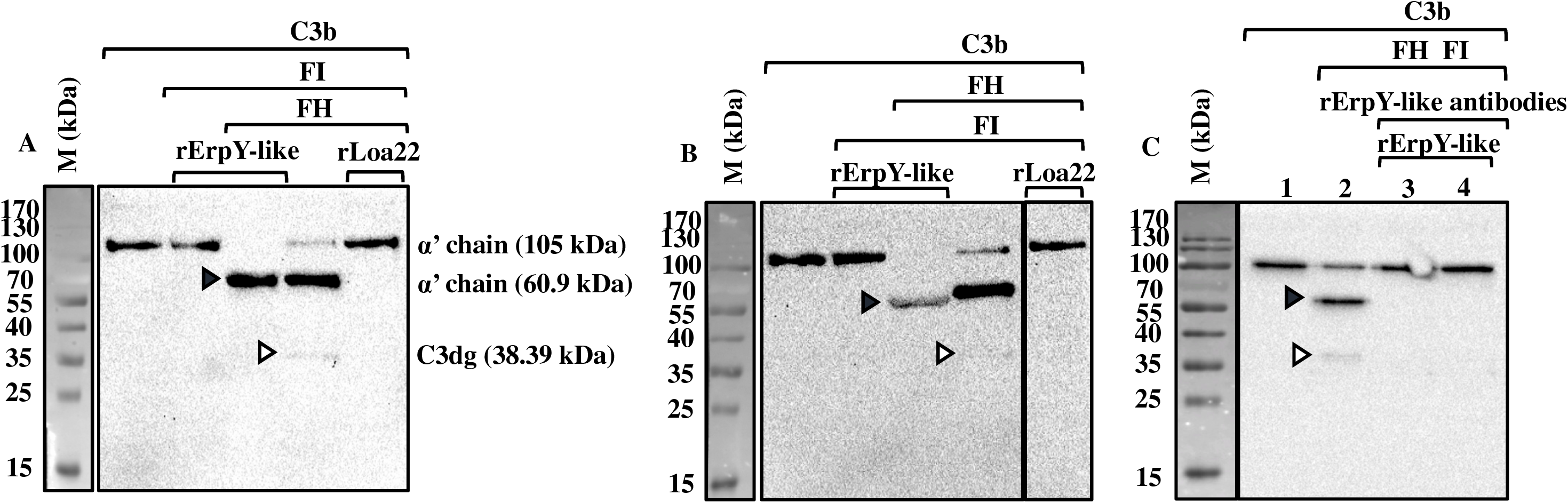
Immunoblot of C3b cleavage assay in the presence of rErpY-like protein. Microtiter plate was coated with either rErpY-like protein or rLoa22 (negative control) and then overlayed with either FH or FI. Unbound components were washed away. The mixture of C3b + FI or C3b + FH were added to the wells initially overlayed with FH and FI, respectively. After 4 h of incubation at 37°C, the supernantant obtained was used for immunobloting with C3b antibodies. **A**. Immunoblot showing FH in the bound state to immobilized rErpY-like protein can cleave C3b. Cleavage of C3b was detected only from the the supernatant of the wells coated with rErpY-like protein or in the positive control, where all three reaction components (FH, C3b, FI) were added together and incubated in BSA blocked wells. **B**. Immunoblot showing FI in a bound state to immobilized rErpY-like protein can cleave C3b. **C**. Immunoblot showing immobilized rErpY-like protein blocked with ErpY-like antibodies cannot bind to FH or FI. Binding of complement regulators to rErpY-like protein was indirectly measured by immunoblotting with C3b antibody. The rErpY-like protein was coated in microtiter plates, blocked with ErpY-like antibodies, and overlayed with FH or FI. C3b + FI or C3b + FH were added to the FH or FI overlayed wells respectively. After 4 h of incubation at 37°C, the supernatant of the FH or FI overlayed wells were resolved in lane 3 and 4 of the 12% SDS polyacrylamide gel, respectively. The C3b cleavage fragments (indicated by triangles) were detected by immunoblotting with a rabbit polyclonal anti-human C3b antibody. Reactions omitting FH served as negative controls. The reactions lacking rErpY-like protein coating but blocked with BSA served as positive controls. Protein size and amount determination were made from intact C3b. Coating of plates with recombinant Loa22 served as a negative control. M depicts the protein standard molecular mass markers.

Previously, our group demonstrated the ability of the rErpY-like protein to bind with FI through ELISA (36). Acquisition of host proteases like plasmin(ogen) by leptospires is an established phenomenon (64), yet there is no testimony about FI acquisition. Thus, we were interested in addressing if the FI bound to rErpY-like protein can also retain its serine protease activity on the C3b. Accordingly, in this case, FI was incubated with the immobilized rErpY-like or rLoa22 on a microtiter plate, and after that, FH in conjunction with C3b was added into the wells. The reaction was accomplished for 4 h, and the supernatant obtained was immunoblotted. The C3b cleavage fragment α’ (60.9 kDa) was detected when FI was pre-incubated with the rErpY-like protein (Fig. 4B). However, no cleavage of C3b could be noticed in the reaction with immobilized Loa22 and when the cofactor was absent.

To further reinforce the explicitness of the C3b cleavage reaction, the immobilized rErpY-like was first blocked with anti-rErpY before the set up of the C3b cleavage assay. Immunoblot of the supernatant of the C3b cleavage reaction did not detect any C3b cleavage product (Fig. 4C). It indicated that blocking rErpY-like protein with specific polyclonal antibodies completely impeded the interaction of the regulatory complement factors (FH and FI) to rErpY-like protein. The antibody-mediated blocking assay also signifies a possible shared interface for the regulatory complement factors (FH and FI) and the specific antibodies towards the rErpY-like protein.

### Recombinant ErpY-like protein inhibits MAC deposition during alternate and classical pathway of complement activation

This study illustrated that regulatory complement factors (FH and FI) that bind to rErpY-like protein could sustain C3b cleavage activity. Moreover, we affirmed that the treatment of bacteria to human serum supplemented with rErpY-like protein results in a gain in viability. Therefore, it was fascinating to address if the gain in bacterial viability during complement-mediated cytotoxicity is due to the reduced MAC deposition, the effector molecule for rupturing bacterial membrane (65). Conditional activation of either the alternative or classical pathways in complement assays was performed *in vitro* utilizing the gelatin veronal buffers (GVB) described elsewhere (66, 67). Activation of CP is Ca^+2^ dependent; thus, GVB supplemented with the divalent Mg^+2^, and Ca^+2^ ions (GVB++) was utilized to selectively initiate the CP whereas, for the alternative pathway GVB with Mg^+2^ and EGTA (a Ca^+2^ ion-selective chelating agent) was utilized (66, 67). The inhibitory effect of the rErpY-like protein on complement activation through one of the pathways (alternate or classical) conditionally was measured using the MAC (C5b-9) deposition levels. It is documented that under *in vitro* condition, zymosan A can act as an initiator of the complement activation for all the three (CP, AP and Lectin) complement pathways (68). Thus, in this study, zymosan A was adopted for activating complement in human serum and was conditionally regulated to a single complement pathway using GVB. A MAC deposition assay was conducted in a microtiter plate where the NHS (20%) diluted with GVB-MgEGTA was used to activate only the alternative pathway. Supplementation of the rErpY-like protein (10-40 µg/mL) to MAC deposition assay resulted in a dose-dependent decline in C5b-9 deposition from 76% to 9%. A ∼90% reduction in MAC deposition was recorded when the NHS was supplemented with rErpY-like (40 µg/mL) versus the NHS without rErpY-like protein supplementation (Fig. 5A). On the contrary, supplementation of rClpP1 or BSA, used as control, did not incite any significant decline in MAC deposition (Fig. 5A). Initiation of complement activation solely through the classical pathway using aggregated human IgG as the activator in NHS (0.5%, diluted in GVB++ buffer) has been reported elsewhere (48, 69). Accordingly, when complement activation assay was mediated solely through the classical pathway, the supplementation of rErpY-like protein (40 µg/mL) resulted in a ∼16% reduction in C5b-9 deposition levels (Fig. 5B). The rErpY-like proteins thus can restrict the MAC deposition stemming from both alternate and classical pathways.

**Figure 5.**
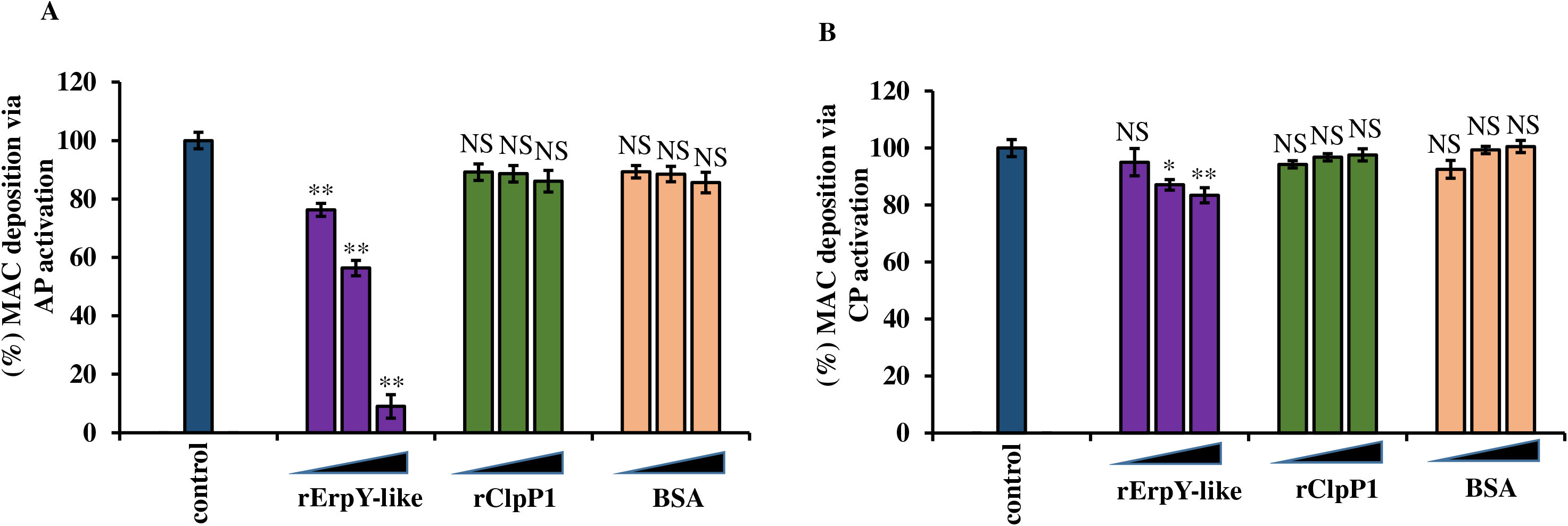
Immunoassay showing rErpY-like protein can inhibit MAC deposition during complement activation. **A**. Reduction of MAC deposition during alternative pathway activation in the presence of rErpY-like protein. Increasing concentrations (10, 20 or 40 µg/mL) of rErpY-like protein, rClpP1 (negative control) and BSA (negative control) was pre-incubated with NHS in GVB MgEGTA, added to microtiter plates coated with zymosan A and incubated for 1 h. **B**. Moderate reduction of MAC deposition during classical pathway activation in the presence of rErpY-like protein. Aggregated human IgG was coated into a microtiter plate. NHS (0.5 % diluted in GVB++) pre-incubated with increasing concentrations (10, 20 or 40 µg/mL) of rErpY-like protein, rClpP1, or BSA was added to the microtiter plate and incubated for 1 h. Activation of the alternative and the classical pathway was probed using rabbit anti-human C5b-9 antibody and its HRPO-conjugated secondary antibodies. The OD values (450 nm) obtained for the reactions were converted to percentage MAC deposition with the OD value for the reaction with only NHS (control) as 100%. The OD values (OD 450 nm) obtained from reactions in NHS supplemented with (rErpY-like, rClpP1, or BSA) were compared with those of NHS (control) using Student’s two-tailed t-test (*, *p* < 0.05, **, *p* < 0,001). The error bars correspond to the respective standard error from three independent experiments performed in triplicates.

### The rErpY like protein activates the complement cascade exclusively through the alternative pathway

By amassing host FH & FI, the rErpY-like protein restricts MAC deposition during the complement activation. Specific outer-membrane proteins of pathogenic bacteria, including streptococcal PepO and SntA (45, 70), have been demonstrated to act as a double edge weapon where it can activate specific pathways of the complement to deplete the complement resources and also thwart the formation of the MAC on the pathogen’s surface (45, 70). Like the ErpY-like lipoprotein, these streptococcal proteins also intervene in the complement activation on the bacterial surface by interacting with the regulatory components of the complement system. Hence, it was interesting to address whether the rErpY-like protein can drain the complement component due to unregulated activation. This may rationalize the decline in MAC deposition in the presence of rErpY-like protein. Therefore, complement activation was executed in diluted NHS (1-6%). NHS was diluted with specific GVB to activate either AP or CP conditionally. The complement protein C3b is a common component in both the alternative and classical pathway. Thus C3b was used to determine the magnitude of complement activation serologically. Zymosan A and aggregated human IgG were utilized to initiate the alternate or classical pathway, respectively, and correlated with that of rErpY-like protein. The C3b deposition levels assessed on immobilized rErpY-like protein due to the activation of the alternative pathway in NHS revealed a similar trend with that of the one activated through zymosan A (Fig. 6A). Interestingly, the rErpY-like protein could not yield C3b when the classical pathway was solely activated versus the aggregated IgG (Fig. 6B). This suggests that rErpY-like protein, besides impeding MAC deposition, can also trigger the alternative path to deplete the host complement components. Thus, ErpY-like protein may aid the spirochetes in shunning the complement-mediated cytotoxicity in the host.

**Figure 6.**
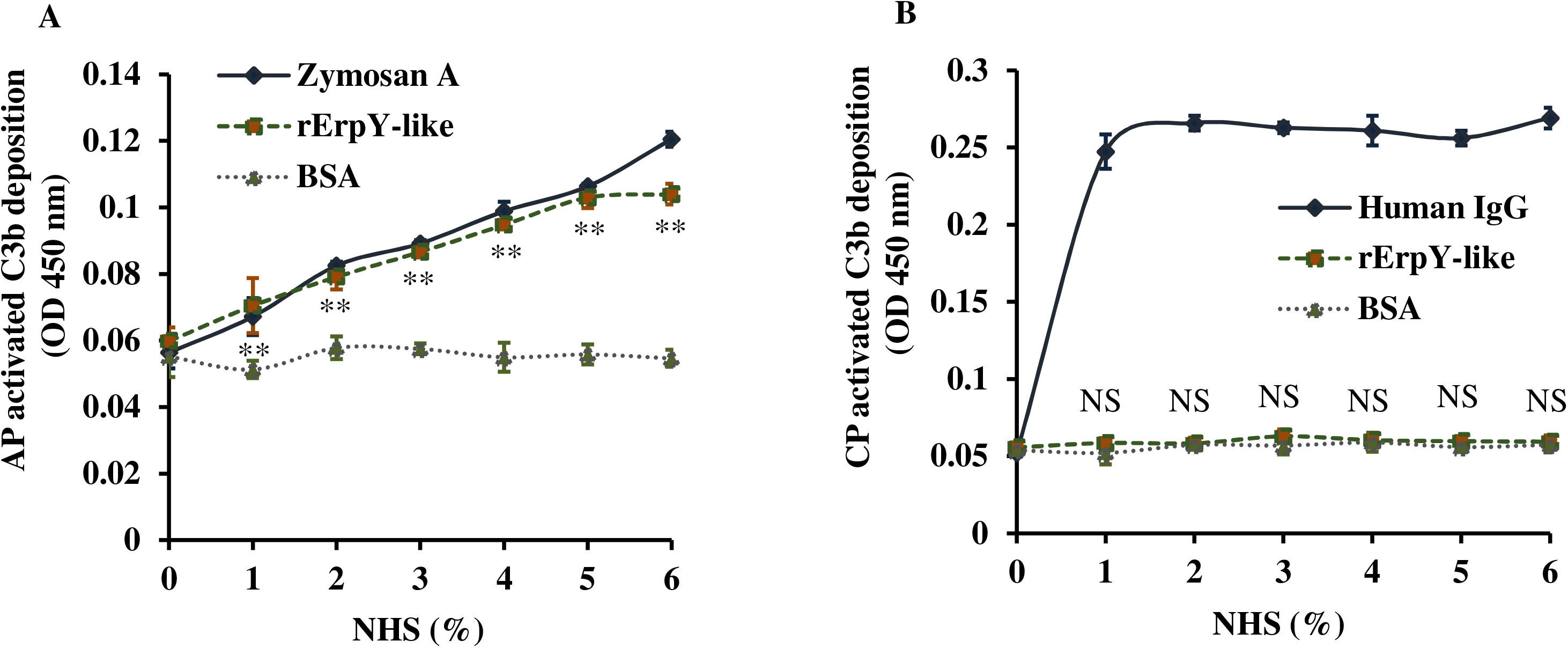
Immunoassay to detect C3b deposition during complement activation in the presence of rErpY-like protein. Microtiter plates were coated with rErpY-like protein, zymosan A (positive control for AP), aggregated human IgG (positive control for CP), or BSA (negative control). The plates were incubated with the indicated concentrations of NHS diluted in GVB MgEGTA for alternative pathway or GVB++ for the classical pathway for 30 min. The detection of C3b deposition using rabbit anti-C3b polyclonal antibodies (1:1000) by ELISA indirectly measured activation of the alternative and the classical complement pathway. **A**. Immunoassay showing C3b deposition during alternative pathway activation by rErpY-like protein. **B**. Immunoassay showing rErpY-like protein cannot activate the classical pathway. Means and standard error of OD values (450 nm) obtained from three independent experiments performed in duplicates are presented. The absorbance values (450 nm) obtained for rErpY-like protein were compared with that of BSA (negative control) using the Student’s two-tailed t-test. (**, *p* < 0.001)

### Specific antibody-mediated blocking of *L. interrogans* surface moderates serum susceptibility

We illustrated the ability of the rErpY-like protein to restrict MAC deposition under *in vitro* conditions. To further validate the role of the ErpY-like lipoprotein of *L. interrogans* in serum resistance, the ErpY-like lipoprotein was blocked with specific antibodies. The complement inactivated anti-rErpY-like mouse sera were utilized for plugging up the ErpY-like lipoprotein on the spirochete. To activate the alternative pathway in human serum, the plugged spirochetes were treated with 20% NHS (diluted in GVB-MgEGTA). The deposition of C5b-9 on the spirochete surface solely through the alternative pathway was measured serologically. A significantly higher level of MAC deposition was detected in the plugged spirochetes versus the controls where the spirochetes were left untreated or pretreated with heat-inactivated mouse preimmune serum (Fig. 7). Our data imply that the ErpY-like lipoprotein of *Leptospira* is involved in host complement immune evasion by restricting the formation of MAC through AP.

**Figure 7.**
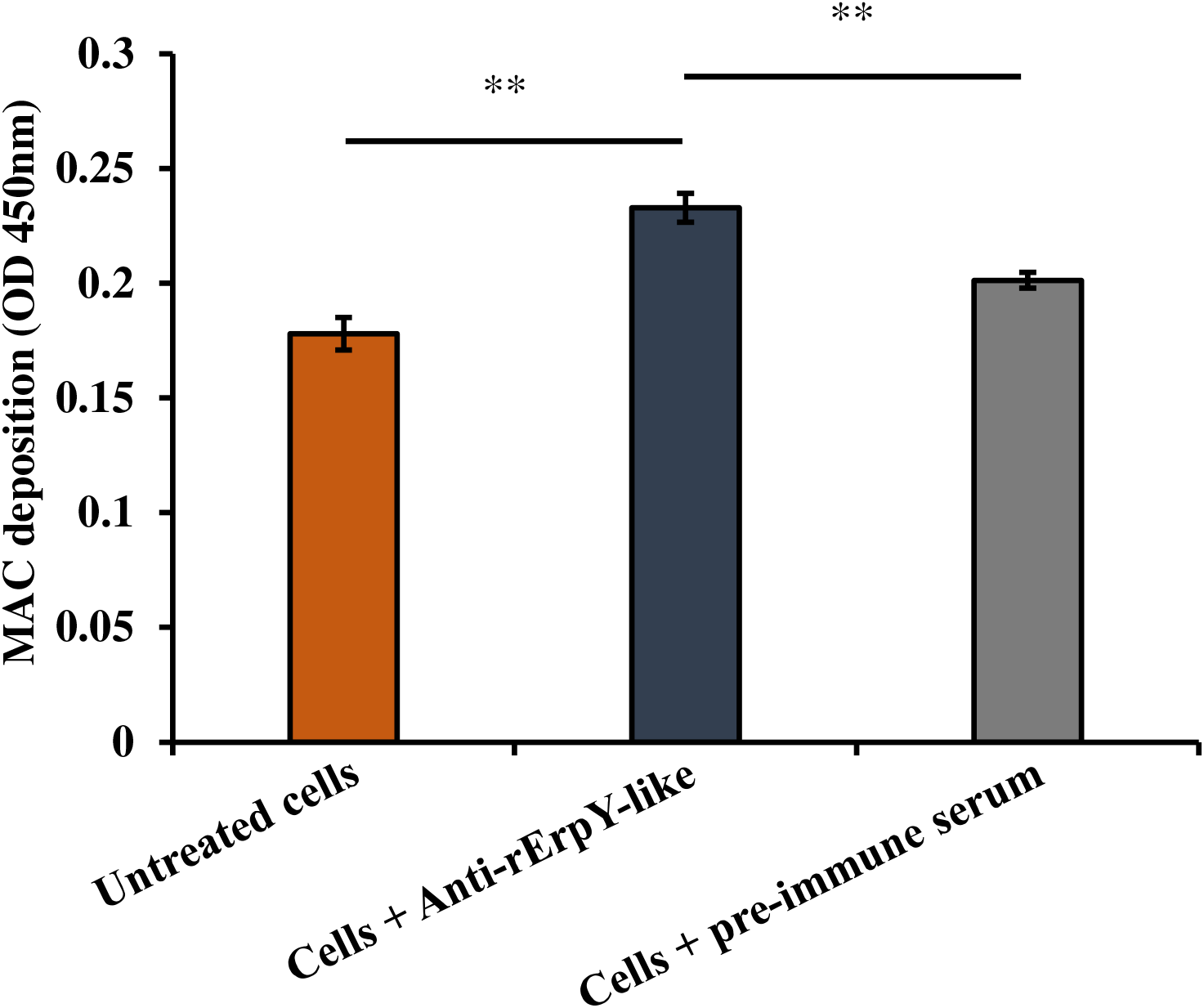
Pathogenic *L. interrogans* blocked with rErpY-like antibodies show increased MAC deposition through the alternative pathway. The deposition of the MAC on the bacterial surface was serologically detected using an anti-human C5b-9 antibody. In this assay, the spirochetes were incubated with complement inactivated anti-rErpY-like or preimmune mouse serum (1: 50) for 1 h, followed by the addition of NHS (20%, diluted in GVB-MgEGTA). The spirochetes were harvested by centrifugation, coated in a microtitre plate and then probed with a C5b-9 antibody (1:1000). The absorbance values (450 nm) obtained for leptospires pre-incubated with rErpY-like antibody were compared with those of cells pre-incubated with preimmune mouse serum and with untreated cells using Student’s two-tailed t-test (** *P* < 0.001). The error bars correspond to the respective standard error from three independent experiments performed in triplicates.

## Discussion

The pathogens must deter host complement activation on their surface and the surrounding microenvironment for thriving, dissemination, and proliferation in the host (Meri, Murgia et al. 2005; Barbosa, Abreu et al. 2009). One of the most widespread strategies embraced by pathogens to elude complement-mediated cytotoxicity is exploiting their surface-expressed proteins to acquire the host complement regulators circulating in the plasma (31, 33). Host synthesize distinct complement regulators like FH, FH-related protein 1, and C4BP to regulate the complement pathway activation in an accountable way and avert any collateral injury to its cell surface while reacting against invaded pathogens (71). Regardless, various microbes, including pathogenic *Leptospira*, have co-evolved with the host and can imitate the host strategy to bypass complement activation. Spirochete *B. burgdorferi* possesses multiple OMPs including CspA (54, 55, 72), CspZ (73), and the Erp family proteins (ErpP, ErpC, and ErpC) (52, 53, 56) that have been distinguished to have a role in complement evasion. Similarly, a list of surface-exposed membrane proteins (LenA, Len B, LcpA, LigA, LigB, and Lsa23) of *Leptospira* can acquire the soluble human complement regulators FH, FH-related protein 1, and/or C4BP (5, 40, 64).

The ErpY-like lipoprotein is one of the recently characterized OMPs of pathogenic *Leptospira* that binds to host ECM components and soluble serum proteins (36). In this study, the immunoelectron micrograph of the spirochete demonstrated that the ErpY-like lipoprotein is surface-exposed and is expressed under the given *in vitro* condition. Our outcomes agree with the prior report where this protein was localized to the bacterial surface using phase separation and proteinase K accessibility assay (36). In the same study, ErpY-like lipoprotein was accounted to bind to complement regulator components (FH and FI). FH binding proteins reckon strategic spatial positioning to nurture efficient evasion of the complement-mediated killing. This motivated us to decipher the role of the ErpY-like lipoprotein of spirochetes in evading complement-mediated destruction. The complement proteins in serum can clear nonpathogenic *L. biflexa* is a well-observed manifestation after activation (8). This study illustrated a significant gain in the viability (%) of bacteria (*E. coli* and *L. biflexa*) when treated with human serum supplemented with rErpY-like protein. In consensus, the supplementation of various known OMPs like Lsa23 in *Leptospira* (Siqueira, Atzingen et al. 2016), PepO in *Streptococcus* (45) and CspA in *Borrelia burgdorferi* (74) enhanced the viability of these microbes in host serum. In pathogenic leptospires, the serum resistance is attributed to the collective endeavor of multiple other OMPs, including LenA, Len B, LcpA, LigA, LigB, Lsa23 (64). Thus in this study, supplementation of rErpY-like protein furnished a moderate serum resistance to the susceptible microorganisms (*E. coli* and *L. biflexa*).

FH is a multimodule (20 CCPs module) glycoprotein, and with the help of such modules, it can bind simultaneously the host cell markers (polyanions) and the soluble C3 fragments to outsmart AP activation on the host surface (61, 75). The N-terminal module of FH (CCPs 1-6) is responsible for the cofactor activity, and also it can bind C3 and C3b complement components (76). CCPs module 13-15 bind C3b, whereas the CCPs 19-20 engage with C3b fragments (iC3b, C3d) (Jokiranta, Hellwage et al. 2000; Haque, Cortes et al. 2020). The regulatory complement FH binds explicitly to host cells (self) expressing polyanions on the cell surface. The polyanionic recognition markers sialic acids (77, 78) is recognized by CCP 20 whereas, the glycosaminoglycans (GAGs) such as heparins (26, 27) and dextran sulfate (25, 27) are recognized CCPs 6-8 (76, 79). These polyanions bound to FH play a leading role in safeguarding the host against unsolicited AP activity on the cell (self) surface. Human-specific pathogens *Neisseria meningitidis* and *N. gonorrhoeae* sequester FH by expressing a host sialic acid analogue on its surface (80-82). Besides expressing the host recognition markers, pathogens including *Leptospira* also express various FH-binding OMPs (LenA, Len B, LcpA, LigA, LigB, Lsa23) that can acquire FH by protein-protein interactions (64). Predominantly these interactions occur through modules excluding the N-terminal region of FH (61, 64), thus, keeping the sequestered FH catalytically active as a cofactor (da Silva, Miragaia Ldos et al. 2015; Siqueira, Atzingen et al. 2016). This study shows that the recombinant ErpY-like protein acquires FH in its active form, as is evident from the cleavage of C3b in the *in vitro* protease assay. Similar instances can be found in the case of leptospiral Lsa23 (38), LcpA (34) and, Lig family proteins (da Silva, Miragaia Ldos et al. 2015). The C3b cleavage assay in the presence of these immobilized leptospiral proteins (Lsa23, LcpA, and Lig family proteins) resulted in cleavage of the α’ chain (109.01 kDa) into multiple fragments (68 kDa, 46 kDa, and 43 kDa) when probed with anti-C3b antibody. In this study, the C3b cleavage products; α’ chain (60.9 kDa) and C3dg (38.39 kDa) fragments were detected when the reaction was performed in the absence of immobilized rErpY-like protein. When the C3b cleavage assay was carried out in the presence of immobilized rErpY-like protein, the C3dg (38.39 kDa) fragment could not be detected. We speculate that the C3dg fragment (38.39 kDa) is sequestered through the CCPs 19-20 module of FH bound to immobilized rErpY-like protein, as described previously (75, 76, 83). On the other hand, α’ chain 2 (37.7 kDa), the first product of FI catalyzed C3b cleavage event, cannot be probed using anti-C3b antibody because anti-C3b used here was generated against the C3 epitope (C3^839-981^), which is not present in α’ chain 2 (37.7 kDa).

To date, the mechanism of interaction between ErpY-like protein and FH is unknown. Nevertheless, the acquisition of FH by the rErpY-like protein in its catalytically active form leads us to speculate that the N-terminal region of FH may remain free. Moreover, rErpY-like protein also plays a role in adhering with multiple host ECM components, including chondroitin sulphates, a polyanionic GAG (36). Notably, both FH and ErpY-like proteins have an affinity with host polyanions. Thus, the ErpY-like lipoprotein binding to the other polyanionic cell markers such as sialic acids, heparins, and dextran sulphate and its competitiveness with FH will be a fascinating topic of a prospective study. Pathogenic leptospires can acquire host proteases (plasminogen/plasmin) from plasma and interfere in the complement activation by degrading the membrane-bound/free complement proteins (C3b, C4b, and C5) (84, 85). Our previous study illustrated that the rErpY-like protein could bind to FI (a serine protease) *in vitro* by indirect ELISA. In this study, FI in a bound state to the rErpY-like protein retains its protease function in coalition with FH and could cleave the C3b molecule. The acquisition of active FI by any OMP of a pathogen has not been documented to date. We speculate that the proximal acquisition of the enzyme (FI) and its cofactor (FH) on the pathogen’s surface helps degrade the surface-bound or the fluid-phase C3b. It may also be an alternate strategy employed by bacteria to evade complement.

It is recounted that pathogenic *Leptospira* can also meddle in complement activation via the CP and LP (Lectin pathway) by sequestering C4b binding protein (C4BP) from human serum on their surfaces (86). C4BP acts as the cofactor for FI for the cleavage of C4b, a complement component constituting the C3-convertase of the classical pathway (87). Multiple leptospiral OMPs (LigA, LigB, LcpA, Lsa23 and, Lsa30) that bind to human FH also shows an affinity for C4BP from serum (34, 43, 88, 89). In contrast, ErpY-like lipoprotein in this study did not demonstrate an affinity for C4BP through indirect ELISA (data not shown).

The recombinant ErpY-like protein displayed a significant inhibitory effect on MAC (C5b-9) deposition mediated either through the alternative or classical pathway. The reduction in the magnitude of MAC deposition through AP in the presence of rErpY-like protein was anticipated because acquiring FH in its active form ends up in a reduction in the C3 convertase of AP. In a recent study, Lsa23, a leptospiral protein, has been proposed to have an inhibitory effect on both AP and CP due to its ability to acquire the catalytically active FH and C4BP proteins, respectively. (38). Likewise, Lig proteins of *Leptospira* spp. can bind various complement regulators (FH, FHL-1, FHR-1 and, C4BP) and hamper both AP and CP activation (90). Among other bacterial species, inhibition of the CP has been recounted in the presence of PepO & SntA from *Streptococcus* spp. that can interact with C1q (45, 70). Unlike others, CspA of *B. burgdorferi* exercises an inhibitory impact on the complement cascade by binding FH and C7 of the alternate and CP, respectively (74). In the presence of rErpY-like protein, a reduction in C5b-9 deposition was detected close to ∼16% through the classical pathway. Nevertheless, the rErpY-like protein did not interact with C4BP, the principal regulator of the classical pathway. Reasonably, the rErpY-like protein may intervene the CP by some other specific regulator (C1-Inh) or the terminal complement components (C7, C8, and C9) like the one reported for leptospiral Lsa23, LIC13259, and LIC12587 (91-93). It is plausible that the inhibition of the complement pathways in the presence of the ErpY-like lipoprotein involves a multilevel mechanism that is yet to be resolved.

In this study, the rErpY-like protein activated the alternative pathway of the complement system. The pronounced inhibitory impact of the rErpY-like protein on MAC deposition and the increased viability of the nonpathogenic bacteria (*E. coli and L. biflexa*) can be attributed to rErpY-like protein’s ability to deplete the serum complement proteins through complement consumption like the streptococcal PepO and SntA (45, 70). Some pathogens like *S. enterica* (62), *Yersinia pestis* (62), *Candida albicans* (94), and *T. denticola* (63, 94) can cleave and inactivate FH. Such inactivation of FH results in unregulated activation of AP and thus depletion of complement constituents in the nearby area. In addition, a shortage of FH may also lead to complement activation on self-cells (host), compromising the tissue integrity, and may enable pathogen dissemination (62, 95). The susceptibility of pathogenic *L. interrogans* to complement was elevated when the spirochetes were blocked with antibodies of rErpY-like protein. This agrees with the previous report where pathogenic leptospires when blocked with Lsa23 antisera, the susceptibility to complement was elevated (38).

Comprehending host-*Leptospira* interaction using the complement regulator FH and FI furnishes insight into the delinquency of regulatory complement component to discriminate self from non-self and the striking resourcefulness of pathogenic leptospires to outmaneuver complement regulation. Analysing *Leptospira* interaction with FH may additionally decipher more evidence about how FH governs in the host. The selective pressure of maintaining FH binding proteins within pathogenic clads of spirochete suggest unnoticed facts about the role of complement in immune defense against pathogens. In sum, this study expands the repertoire of proteins associated with the development of complement resistant phenotype in pathogenic *Leptospira*. ErpY-like lipoprotein sequesters active FH and FI onto the bacterial surface thus, downregulates the alternative pathway. The recombinant ErpY like protein activates the alternative complement pathway, which may deplete the complement protein in its microenvironment and inhibit MAC formation.

## Acknowledgements

The authors gratefully acknowledge ICMR, Port Blair India, for providing the *Leptospira* strains. We acknowledge the central instrumentation facility (CIF), IIT Guwahati, for electron microscopy. We would also like to thank the department of microbiology, College of Veterinary Science, Guwahati, for generating polyclonal antibodies.

